# Optimization of crossing strategy based on the usefulness criterion in inter-population crosses considering different genetic effects among populations

**DOI:** 10.1101/2025.01.21.634020

**Authors:** Sei Kinoshita, Kengo Sakurai, Kosuke Hamazaki, Takahiro Tsusaka, Miki Sakurai, Kenta Shirasawa, Sachiko Isobe, Hiroyoshi Iwata

## Abstract

In the breeding programs of self-pollinated plants, achieving genetic improvement in multiple traits can be challenging when relying solely on a single biparental population. Interpopulation crosses are employed to integrate favorable alleles from multiple biparental populations to overcome this limitation. In this context, it is crucial to consider the distinct genetic effects in different populations. In this study, we utilized a selection method based on the usefulness criterion (UC) to identify cross pairs suitable for interpopulation crosses. We expanded this approach to enhance breeding programs accounting for varying genetic backgrounds within the genomic selection framework. Using the medicinal plant red perilla as the study material, we conducted simulations to compare the efficacy of selection based on estimated genomic breeding values with that based on UC. Our findings demonstrate that the proposed method is effective in facilitating the simultaneous improvement of multiple traits, particularly by considerably increasing genetic gains among the top-performing individuals in the population. Furthermore, we provide guidelines for implementing interpopulation crosses, including recommendations for the optimal generation for crossing and the appropriate reference generation for calculating the UC. The results obtained in this study offer valuable insights for small-scale breeding programs aimed at simultaneously enhancing multiple traits through inter-population crosses and are applicable to a wide range of crops, including neglected and underutilized species.

**Key Message:** Herein, a method has been proposed for selecting optimal cross pairs based on the genetic potential of progeny in inter-population crosses, considering different genetic effects among populations.

## Introduction

Genomic selection (GS) utilizes genome-wide marker information to predict the genetic potential of individuals and selects them based on the predicted genotypic values (Meuwissen et al. 2001). Initially, GS was used in dairy cattle breeding (Hayes et al. 2009); however, more recently, it has been actively used in breeding various plant species (Bernardo and Yu 2007; Yamamoto et al. 2016; Mahadevaiah et al. 2021). GS has helped develop several breeding schemes tailored to the specific traits of different plant species. For wheat breeding, recurrent selection within a single biparental population is commonly employed (Bassi et al. 2016). The simulation studies of wheat breeding programs have typically incorporated strategies such as random mating, selfing, and the production of doubled haploids (DHs) (Daetwyler et al. 2015; Gaynor et al. 2017). However, these studies generally focused on repeated mating or selfing within a single biparental population, with few considering using multiple parents as the initial crossing parents in the context of GS.

Many breeding programs throughout the history of plant breeding have used multiparental populations. Using multiple parents is believed to increase genetic diversity and facilitate the development of varieties with superior traits (Arrones et al. 2020). Studies have focused on integrating multiple desirable genes into a single population by crossing different populations derived from various parental lines in a process known as gene pyramiding (Ramalingam et al. 2020). In genomic breeding using multiple parents or populations, it is crucial to consider the distinct genetic effects of each parent or population. Models for quantitative trait locus (QTL) analysis suited to multiple populations with different genetic backgrounds have been developed, assuming that the effects and locations of QTLs may vary (Li et al. 2021). These models have been applied to various plants and are considered to enhance the detection power of QTLs (Mangino et al. 2022).

Distinct genetic effects within different populations are particularly advantageous for small-scale breeding programs. For example, the genetic traits of neglected underutilized species (NUS), which have recently gained attention because of their high nutritional value and adaptability to sustainable agriculture, can be improved using efficient breeding strategies, such as GS (Ye and Fan 2021). However, it is challenging to implement large-scale breeding programs for NUS because of the limited genetic and economic resources available for such species. In such cases, a practical approach involves selecting a promising set of lines, creating multiple biparental populations, and subsequently combining these populations. It has been hypothesized that in such small-scale breeding populations, allele effects vary based on population (Maurer et al. 2017). In this study, we focused on the medicinal plant red perilla, a type of NUS that has undergone small-scale breeding in multiple biparental populations. These perilla populations exhibit superior characteristics, as they produce various bioactive compounds with medicinal value; however, there remains a pressing need to simultaneously improve the contents of multiple bioactive compounds in these populations (Kinoshita et al. 2023). In this study, we investigated multiple biparental populations with distinct genetic effects in the context of GS to evaluate the effectiveness of inter-population crosses for enhancing multiple traits.

In plant breeding programs, new varieties are developed through repeated crossing and selection cycles. Thus, it is essential not only to select superior individuals but also to identify optimal mating pairs that can produce high-performing progeny. Typically, mating individuals with high genotypic values ensures high performance in the population mean of the progeny. However, in breeding, it is crucial to maintain not only a high progeny mean but also the ability to produce superior progeny over the long term (Sanchez et al. 2023). Selecting the cross pairs capable of generating progeny with high genetic variance is necessary to sustain genetic gains over time.

In 1975, Schnell and Utz introduced the “usefulness criterion” (UC) as a measure of the trait mean of the upper fraction of progeny from a given cross. UC is defined as UC = *μ* + *ihσ*, where μ is the mean breeding value of parents, *i* is the selection intensity, *h* is the square root of heritability, and *σ* is the square root of the genetic variance in the progeny. With the advent of high-density marker information, integrating genomic prediction (GP) with UC has become feasible, enabling the prediction of genetic variance in progeny using genomic estimated breeding values (GEBVs). Initially, the genetic variance of the progeny was predicted by simulating their genomes (Iwata et al. 2013; Bernardo 2014; Mohammadi et al. 2015). However, as the number of potential crosses and progeny increase, the computational cost of the simulations becomes prohibitive. To address this issue, Lehermeier et al. (2017) proposed an analytical approach to calculate the genetic variance of progeny using estimated marker effects and recombination rates. Further advancements have expanded this method beyond simple biparental crosses, improving the computational efficiency for more complex breeding designs such as three- and four-way crosses (Allier et al. 2019; Danguy des Déserts et al. 2023). In these studies, breeding programs typically focused on developing recombinant inbred lines (RILs) or DHs through biparental crosses, with no examples of their application to interpopulation crosses.

In the present study, we integrated the methods developed by Lehermeier et al. and Danguy des Déserts et al. to compute the genetic variance of progeny derived from crosses between multiple biparental populations with different marker effects. The computed genetic variance was then used as a criterion to select the cross pairs. In this study, we aimed to evaluate whether interpopulation crosses between populations with different genetic effects can effectively and simultaneously improve multiple traits in the context of GS. In addition, we sought to propose guidelines for conducting such interpopulation crosses. Therefore, we performed simulation-based analyses using data from two red perilla RIL populations, each of which excels in different traits. Specifically, we propose a method for selecting mating pairs using UC, which accounts for marker effects varying across populations. Additionally, we reveal the optimal generation at which these crosses must be performed to achieve desired outcomes. Our findings provide valuable insights for guiding small-scale breeding programs aimed at simultaneously enhancing multiple traits through inter-population crosses and can be applied to a wide range of crops, including NUS.

## Materials and methods

### Plant materials

The plant materials used in this study consisted of two breeding populations of the F_4_ generation from two-way crosses between red and green perilla (Kinoshita et al. 2023). These populations were generated by crossing ‘SekihoS8’ × st27 (S827) and ‘SekihoS8’ × st40 (S840). Hereafter, these populations will be referred to as S827 and S840, respectively. ‘SekihoS8’ is a representative variety of red perilla (*Perilla frutescens var. crispa f. purpurea*) provided by TSUMURA & CO, Japan. Moreover, st27 and st40 are green perilla (*P. frutescens Britton var. crispa* Decne.) lines obtained from germplasm collections managed by the Genebank of the National Agriculture and Food Research Organization, Japan. The S827 and S840 populations had 298 and 297 individuals with different genotypes, respectively. Further details of these two populations have been described by Kinoshita et al. (2023).

### Reference genome construction

The DNA of ‘SekihoS8’ was extracted using the Genomic-tip kit (QIAGEN Technologies). Genome sequencing was performed using the Sequel II platform (Pacific Biosciences), and HiFi reads were generated with the CCS version 4.2.0 (Pacific Biosciences) software, requiring a minimum of three subreads per read. Genome assembly was performed using hifiasm v0.16.1 (Cheng et al. 2021), and candidate chloroplast and mitochondrial genome sequences were eliminated using the chloroplast genome sequence of *P. frutescens* (NC_030755.1) and the mitochondrial genome sequences of *Scutellaria galericulata* and *Ballota nigra* (OX335799.1, OX344731.1, OX344732.1, and OX344733.1) as reference sequences. The resulting scaffolds were aligned to the previously reported genome sequence of perilla cultivar ‘Hoko-3’ (Tamura et al. 2023) using the RaGOO tool (Alonge et al. 2019). Genome structure comparisons were performed using D-Genies (Cabanettes and Klopp 2018). Genome structure analyses using k-mers and telomere sequence detection were performed using Smudgeplot and tidk, respectively (Ranallo-Benavidez et al. 2020; De la Rosa and Mark 2023). Gene predictions were performed using Helixer 0.3.2 (land_plant_v0.3_a_0080. h5) (Stiehler et al. 2021). The accuracy of the assembled genome and predicted gene sequences was evaluated using the benchmarking universal single-copy orthologs (BUSCO) v5.2.2 (obd10) tool (Simão et al. 2015).

### Genotype and phenotype data

The F_4_ generations, S827 and S840, were phenotyped for two traits: perillaldehyde and rosmarinic acid contents. These are the main medicinal compounds in perilla and are widely used in traditional herbal medicine. A detailed description of the methods for measuring perillaldehyde and rosmarinic acid contents has been provided earlier (Kinoshita et al. 2023).

All 595 individuals from the F_4_ generations of S827 and S840 were genotyped using the double-digest restriction site–associated DNA sequencing (dd-RAD-Seq) method (Shirasawa et al. 2016). The sequencing of the dd-RAD-Seq library was performed using a DNBSeq-G400RS (MGI Tech Co. Ltd., Shenzhen, China), generating 100 bp reads. The obtained reads were then mapped onto the assembled sequence of ‘SekihoS8’ using Bowtie2 (Langmead and Salzberg 2012), and variant calling was performed with the bcftools in SAMtools (Li et al. 2009). After quality control and filtering based on minor allele frequency, as described in Kinoshita et al. (2023), 1,951 single nucleotide polymorphisms (SNPs) were obtained across the entire population, with 692 SNPs in S827 and 1,439 SNPs in S840. The genotype scores were coded as 0 for homozygous SNPs matching ‘SekihoS8,’ 1 for heterozygous SNPs, and 2 for homozygous SNPs matching the other parents.

### GP model

The BayesB model (Meuwissen et al. 2001) was used for GP to estimate marker effects for each trait and population. The BayesB model can be represented using Equation (1) as follows:

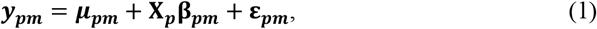

where *N* is the number of individuals in one population, *L* is the number of markers, ***y***_*pm*_ is an *N* × 1 vector representing phenotypic values for *m*^th^ trait in *p*^th^ population, μ_*pm*_ is the overall mean, **X**_*p*_ is an *N* × *L* matrix of marker genotypes for *p*^th^ population, **β**_*pm*_ is an *L* × 1 vector corresponding to marker effects for *m*^th^ trait and *p*^th^ population, and 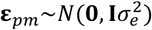 is an *N* × 1 vector of errors, where 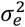 is the error variance. The Markov Chain Monte Carlo was run for 60,000 iterations, with the first 12,000 samples discarded as burn-in, and a sampling interval (thinning) of 5. The BayesB model was implemented using the “BGLR” function in the “BGLR” package version 1.1.0 in R (Pérez and De Los Campos 2014). The estimated marker effects are shown in Fig. 1 and were used in the following simulation studies.

**Fig. 1.**
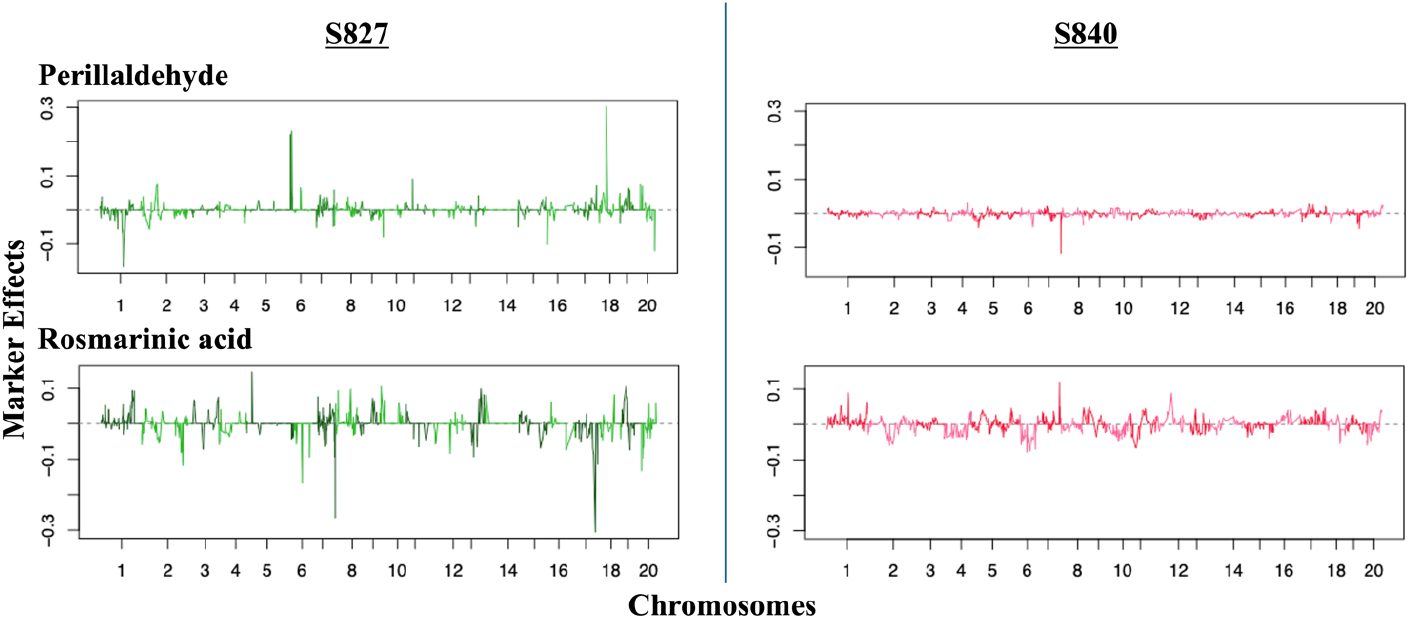
Estimated marker effects of each population and trait. The left and right panels show the marker effects in two biparental populations: S827 and S840, respectively

### Simulations

#### Breeding program

Stochastic simulations of breeding programs were performed to evaluate the effectiveness of interpopulation crosses for multi-trait improvement. An outline of the breeding program is shown in Fig. 2. Two scenarios are investigated in this study. In both scenarios, the first round of selection and crossing was performed at the F_4_ generation (current generation), and a GP model was constructed. The second round of selection and crossing was implemented in G_t_ generation (*t* = 1, 2, 3, 4), which refers to the *t*^th^ generation after the initial cross (i.e., the first round of selection and crossing). In other generations with no selection or crossing, all individuals underwent self-pollination, and the next generation was obtained through single-seed descent (SSD).

**Fig. 2.**
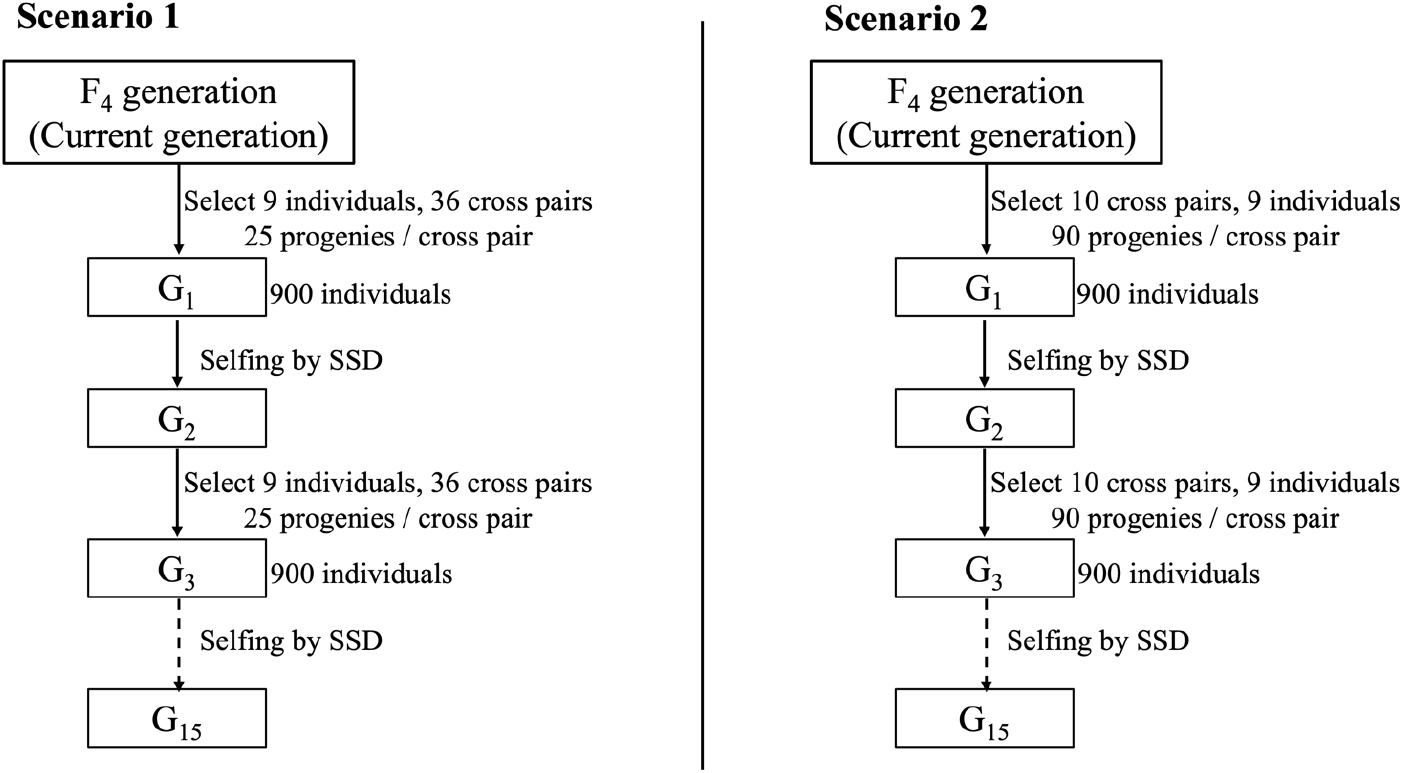
Breeding program used in this study. SSD, single-seed descent

In Scenario 1, high-performance individuals with high genotypic values were selected for crossing. In Scenario 2, cross pairs that could generate superior progeny were selected by predicting the genotypic value of their progeny. The selection of cross pairs in Scenario 2 was based on the predicted genotypic value of the generation used for the second-time cross (G_t_) or final generation (G_15_). The breeding program was designed to span 15 generations to ensure genetic fixation. The details of scenarios 1 and 2 are described in the following section. The simulations were repeated 100 times for each scenario, and the genetic gains were averaged across repetitions to compare different scenarios.

#### Scenario 1: individual-based selection

In Scenario 1, nine individuals (five from the S827 population and four from the S840 population) were selected from the F_4_ generation based on the highest selection index, which was defined as the sum of the GEBVs for the two traits: perillaldehyde and rosmarinic acid contents. The index was calculated using Equation (2) as follows:

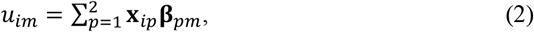

where ***u***_*im*_ represents the scaled GEBVs of individual *i* (*i* = 1, …, 595) for *m*^th^ trait; **x**_*ip*_ is an 1 × *L* vector corresponding to the marker genotype of individual *i* belongs to the *p*^th^ population; **β**_*pm*_ denotes the estimated marker effects of *m*^th^ trait and *p*^th^ population; and *L* is the total number of markers (*L* = 1,951). The selected nine individuals were crossed with each other in all possible combinations without selfing, resulting in 36 combinations. A total of 25 progenies produced from each cross were collected, maintaining a population size of 900. At the G_t_ generation, selection was conducted in the same manner as for the first cross. Nine individuals with the highest selection index were selected and crossed again in all combinations to generate the next generation of progeny.

#### Scenario 2: cross-pair-based selection

In Scenario 2, 10 cross pairs, including nine individuals from the F_4_ generation, were selected based on the UC. UC was used to evaluate the potential of a cross to generate superior progeny and was calculated using Equation (3) as follows:

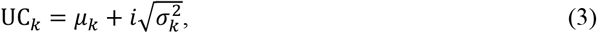

where μ_*k*_ is the mean genotypic value of cross *k, i* is the selection intensity (set to 1.96 in this study), and 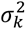 is the genetic variance of progeny generated from cross *k*. The genetic variance, 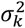, can be calculated analytically for any generation of inbred lines. For this study, we combined the methodologies outlined by Allier et al. (2019) and Danguy des Déserts et al. (2023) to apply the calculation of 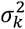 to inter-population crosses, assuming different genetic effects among populations. 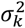 was computed using Equation (4) as follows:

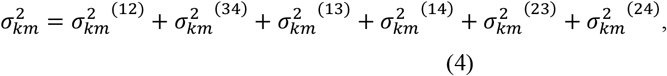

where allele 1 and allele 2 are derived from parent 1, and allele 3 and allele 4 are derived from parent 2. 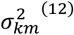 represents the genetic variance between allele 1 and allele 2 from parent 1 for *m*^th^ trait,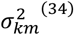 represents the genetic variance between allele 3 and allele 4 from parent 2 for *m*^th^ trait. The terms 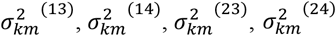 represent the genetic variances between two alleles from different parents. The genetic variance 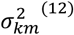 can be computed using Equation (5) as follows:

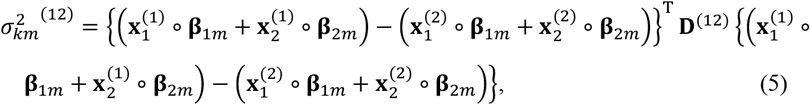

Where 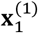 and 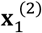 are *L* × 1 vectors representing the haplotype score of allele 1 and allele 2 (coded as 0 or 1) for parent 1, respectively; **β**_1*m*_ is an 1 × *L* vector of marker effects from population 1 for trait 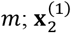 and 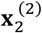 are *L* × 1 vectors representing the haplotype score of allele 1 and allele 2 for parent 1, respectively; **β**_2*m*_ is an 1 × *L* vector of marker effects from population 2 for trait 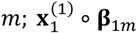 represents the Hadamard product of two vectors; and **D**^(12)^ is an *L* × *L* variance-covariance matrix of linkage disequilibrium between allele 1 and allele 2, which is common for all crosses. The genetic variance 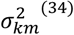, between the two alleles from parent 2, is computed in the same manner. The genetic variance between alleles from different parents, such as 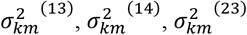, and 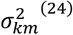 can also be calculated. For example, 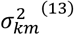can be calculated using Equation (6) as follows:

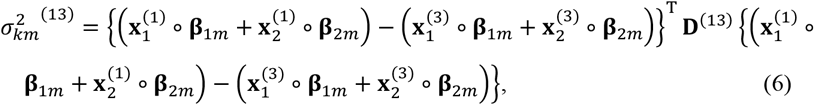

Where 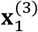 represents an *L* × 1 vector of the haplotype score of allele 3 from parent 2; **β**_1*m*_ and **β**_2*m*_ are 1 × *L* vectors of marker effects from population 1 and 2 for trait *m*. The linkage disequilibrium matrix **D** can be computed based on the recombination rate using Equations (7) and (8) as follows:

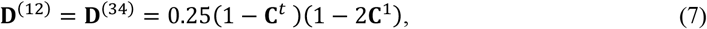

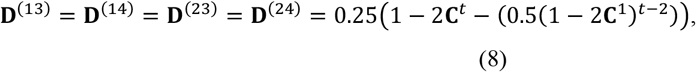

Where 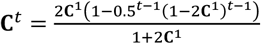 is an *L* × *L* matrix of recombination rates in the *t*^th^ generation (*t* ≠ 1), and **C**^1^ is an *L* × *L* matrix of recombination rates between markers. The diagonal elements of **D** are 0.25 when *t* = 1 and 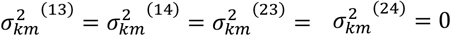.

In this scenario, cross pairs with the highest selection index, defined as the sum of the UC of two traits 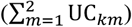, were selected. Here, μ_*km*_, representing the mean genotypic value of cross *k* for trait *m*, were scaled using the mean and standard deviation of genotypic values from the F_4_ generation. Similarly, 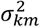, representing the genetic variance of progeny generated from cross *k* for trait *m*, were scaled only using the standard deviation of the genotypic value from the F_4_ generation. After selecting cross pairs based on UC, 90 progenies produced from each cross were collected to maintain a total population of 900. At the G_t_ generation, cross pairs were selected in the same manner as for the first cross. However, unlike that for the first cross, to reduce computation time, UC was calculated for all possible combinations of 450 individuals with the highest selection indices, and 10 cross pairs with the highest UC were subsequently selected to generate progeny.

### Comparison

We conducted 100 breeding simulations for the two scenarios. For comparison, we computed the mean genetic gains of the entire population and the top 1% of individuals in each generation using Equation (9) as follows:

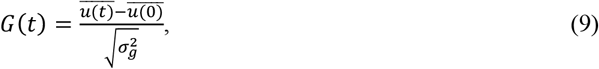

Where 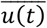 is the mean value of the selection index or GEBV for a trait in the G_t_ (*t* = 1, 2, …, 15) generation for the entire population or top 1% of individuals, 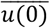 is the corresponding mean for the initial F_4_ population, and 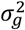 is the genetic variance of individuals in the initial F_4_ population.

We also computed the genetic variance of each generation using Equation (10) as follows:

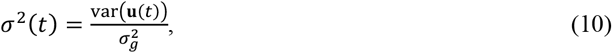

where varb (**u**(*t*)) is the genetic variance of the individuals in the G_t_ generation, and 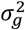 is the genetic variance of the individuals in the initial F_4_ population.

## Results

### ‘SekihoS8’ reference genome construction

A total of 2,011,482 HiFi reads with a length of 36.4 Gb were obtained and assembled using hifiasm v0.16.1 to construct a primary assembly and two haplotype assemblies (Hap1 and Hap2). After removing the mitochondrial and chloroplast genomes, there were 108, 121, and 106 assembled sequences for the primary assembly, Hap1, and Hap2, respectively (Table S1). The total lengths of the assemblies ranged from 1,231 Mb to 1,259 Mb, with N50 values ranging from 55.4 Mb to 56.0 Mb. When aligned to the ‘Hoko-3’ genome, all contigs from these assemblies were successfully mapped to 20 chromosomal sequences, with no major structural variations detected (Fig. S1). The results of the Smudgeplot analysis suggested that ‘SekihoS8’ is a potential allotetraploid (Fig. S2). Additionally, telomere sequences were detected in the terminal regions of most chromosome-scale scaffold sequences (Fig. S3). The constructed genomes were designated as Pfru_SekihoUp_1.0 (primary), Pfru_SekihoH1_1.0 (hap1), and Pfru_SekihoH2_1.0 (hap2). The BUSCO analysis revealed that the percentage of complete genes in all three genome assemblies was 99.4%, indicating the high completeness and coverage of known genes. Gene prediction using a helixer identified 65,161 genes in the primary assembly: 65,078 in Hap1 and 62,784 in Hap2 (Table S2). The results of the BUSCO analysis further confirmed high accuracy, with complete BUSCO scores ranging from 98.7 to 99.1%. Duplicate complete BUSCOs showed the proportions of 89.0% for Hap2 and 99.1% for primary and Hap1 assemblies, indicating a high level of genome duplication.

### Changes in genetic gains for selection index

Changes in the mean genetic gains of the entire population and that of the top 1% of individuals from the initial population (F_4_ generation) to the final generation (G_15_) are shown in Fig. 3. Regarding the mean genetic gains of the entire population, Scenario 2 showed greater improvement than that observed in Scenario 1 in almost all cases, except when a second-round cross was performed between the individuals of the G_1_ generation. In terms of mean genetic gains of the top 1% of individuals, Scenario 2 considerably outperformed Scenario 1 in all cases. The mean genetic gains of the entire population and the top 1% of individuals were higher when the second-round cross was made at a later generation, although the extent of improvement diminished as the generation progressed. Fig. 4 shows the change in genetic gains when the selection of the first-round cross-pairs in Scenario 2 was conducted based on different generations of progeny. Both the mean genetic gain of the entire population and that of the top 1% of individuals were slightly higher when the cross pairs for the first-round cross were selected based on the predicted genotypic value of the generation used for the second-round cross rather than based on the genotypic values of the individuals in the cross pairs.

**Fig. 3.**
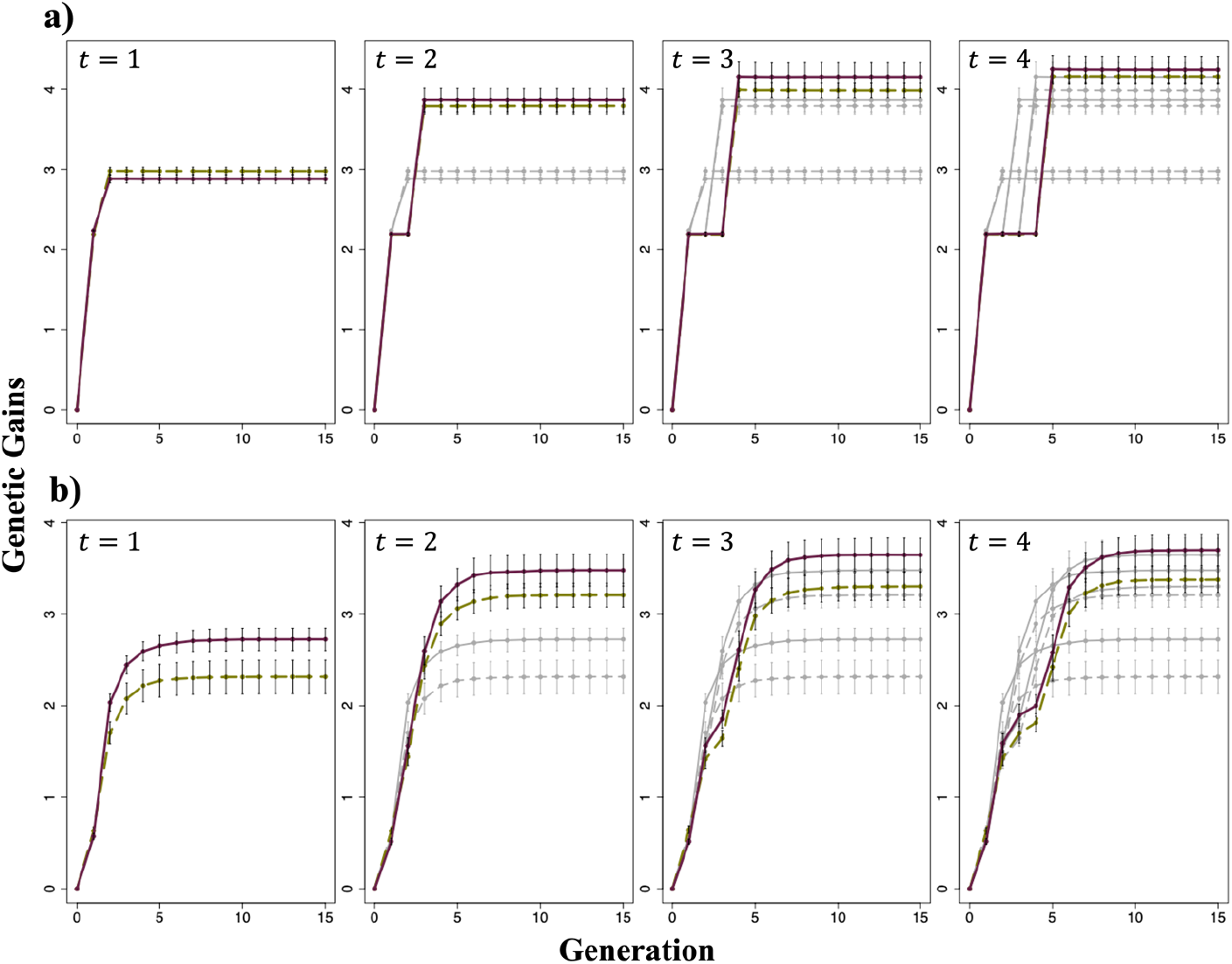
Changes in the mean genetic gains of the entire population and top 1% of individuals evaluated for selection index. (**a**) Changes in the mean genetic gains of the entire population. (**b**) Change in the mean genetic gains of the top 1% of individuals. *t* represents the generation at which the second-round cross was made. Green dashed line: Scenario 1; Purple solid line: Scenario 2

**Fig. 4.**
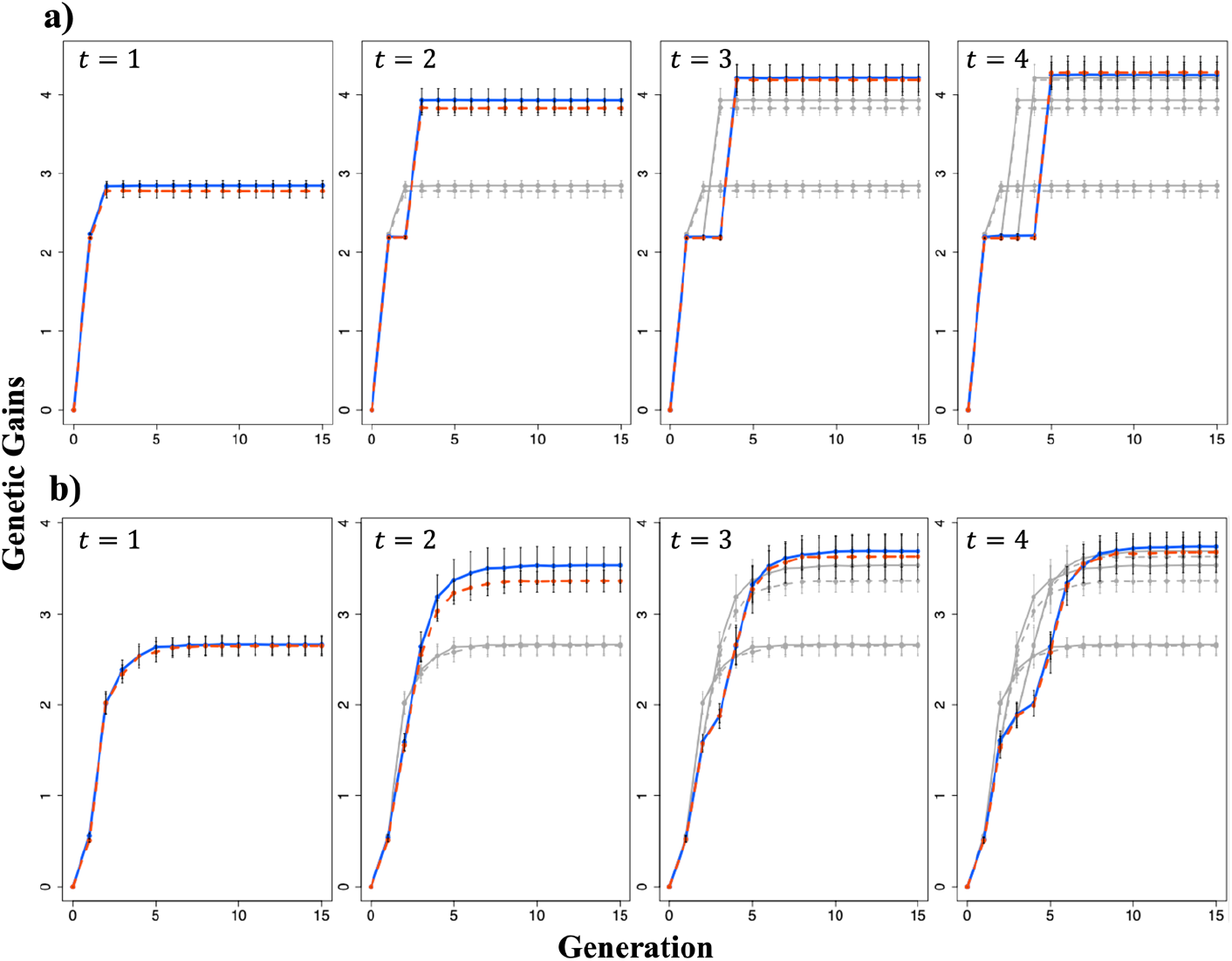
Changes in the mean genetic gains of the entire population and top 1% of individuals when the selection of the cross pairs for the first-round cross was based on the predicted genotypic values of the different generations of progenies in Scenario 2. (**a**) Changes in the mean genetic gains of the entire population. (**b**) Changes in the mean genetic gains of the top 1% of individuals. *t* represents the generation at which the second-round cross was made. Red dashed line: when the cross pairs for the first-round cross were selected based on the predicted genotypic value of the final generation; Blue solid line: when the cross pairs for the first cross were selected based on the predicted genotypic value of the generation used for the second-round cross

### Changes in genetic variance

Fig. 5 shows changes in genetic variance. Scenario 2 maintained a higher genetic variance across all cases. The timing of the second-round cross impacted genetic variance, with later crosses resulting in reduced genetic variance in the final generation (G_15_) but increased genetic variance in the generation at which the second-round cross was made (G_t_, where *t* = 1, 2, 3, 4).

**Fig. 5.**
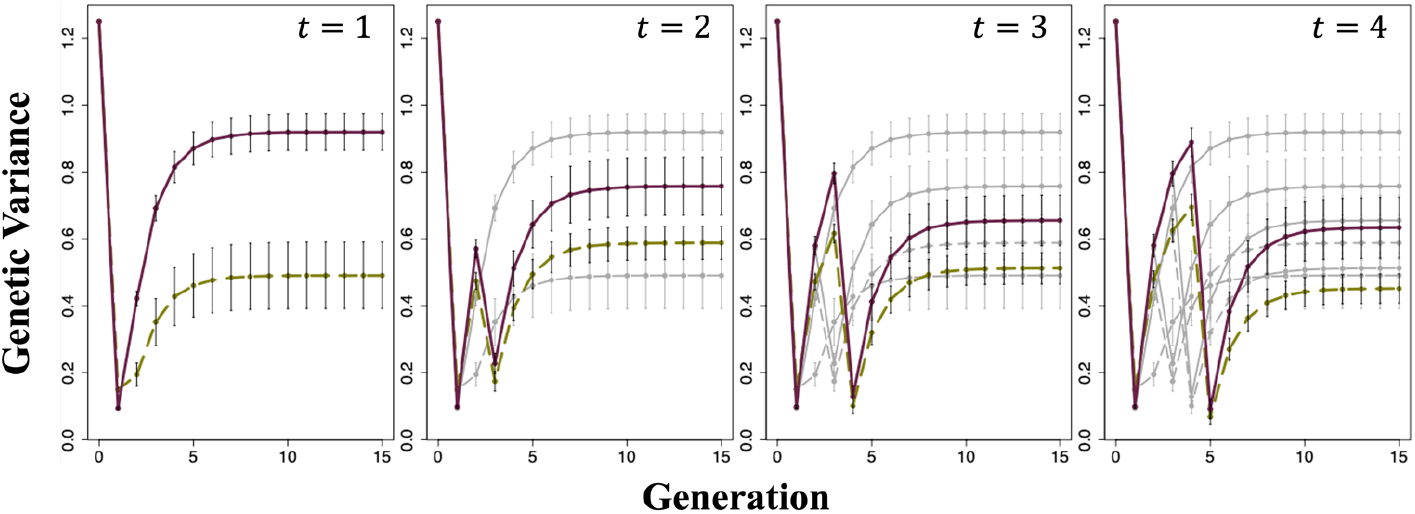
Changes in the genetic variance of the selection index. *t* represents the generation at which the second-round cross was made. Green dashed line: Scenario 1; Purple solid line: Scenario 2

Fig. 6 shows the change in genetic variance when the selection of first-round cross pairs was based on different generations of progeny in Scenario 2. The genetic variance was nearly the same in the generation used for the second-round cross but higher in G_15_ when the cross pairs for the first-round cross were selected based on the predicted genotypic values of G_15_ rather than the genotypic values of the generation used for the second-round cross. However, the opposite trend was observed when the second-round cross was made at the G_1_ generation.

**Fig. 6.**
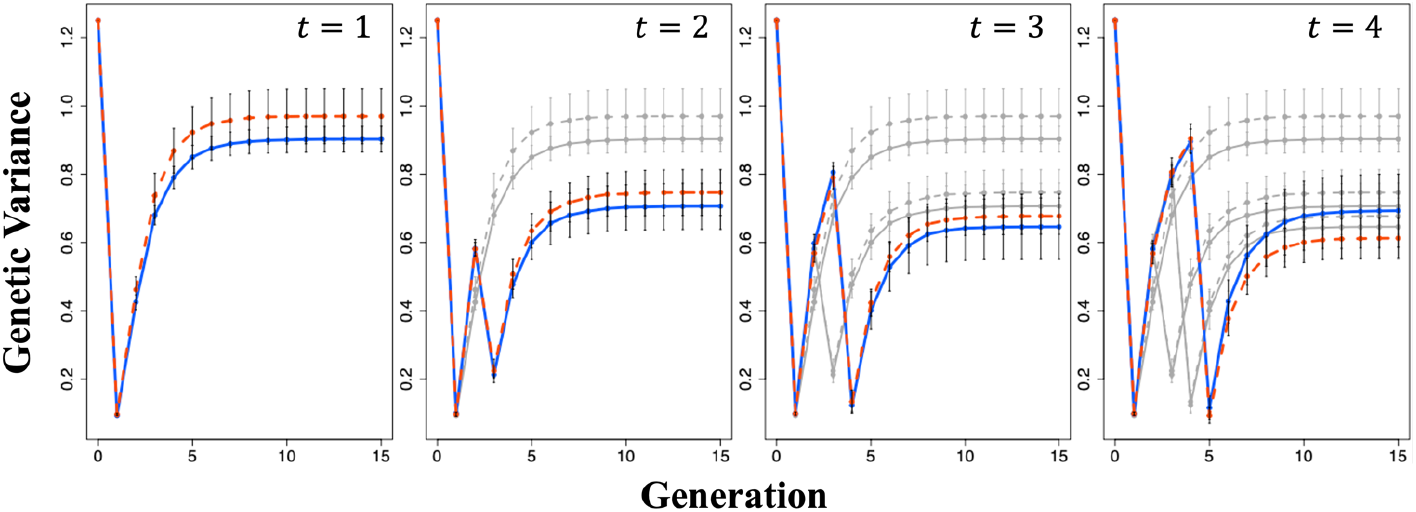
Changes in genetic variance when the selection of the cross pairs for the first-round cross was based on the predicted genotypic values of the different generations of progeny in Scenario 2. *t* represents the generation at which the second-round cross was made. Red dashed line: when the cross pairs for the first-round cross were selected based on the predicted genotypic values of the final generation; Blue solid line: when the cross pairs for the first-round cross were selected based on the predicted genotypic values of the generation used for the second-round cross

### Changes in genetic gains and genetic variance for each analyzed trait

Fig. 7 shows the change in genetic gains of the top individuals and the genetic variance for each trait, namely, perillaldehyde and rosmarinic acid contents. Regarding the mean genetic gains of the top 1% of individuals, Scenario 1 resulted in higher genetic improvement for perillaldehyde content, whereas Scenario 2 led to higher genetic improvement for rosmarinic acid content. Regarding genetic variance, Scenario 2 exhibited higher variance for both perillaldehyde and rosmarinic acid contents than in Scenario 1 at the final generation (G_15_) and lower variance in the generation used for the second-round cross. Genetic variance in the final generation and the generation used for the second-round cross varied depending on the timing of the second-round cross. The descending order of genetic variance in the final generation for Scenarios 1 and 2 was G_2_>G_3_>G_4_>G_1_ and G_1_>G_2_>G_3_>G_4_, respectively. Similarly, the descending order of genetic variance in the generation used for the second-round cross for Scenarios 1 and 2 was G_4_>G_3_>G_2_>G_1_.

**Fig. 7.**
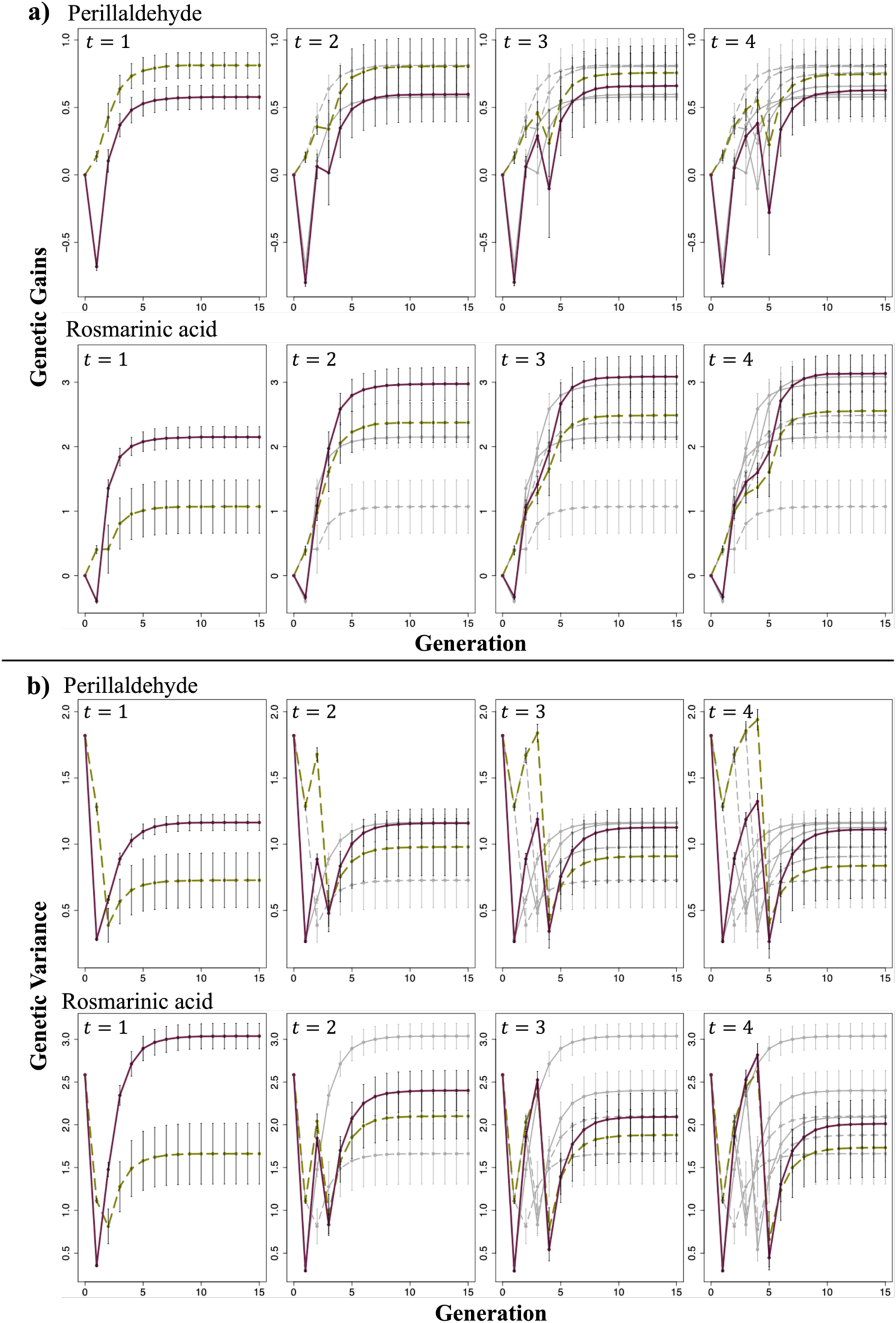
Changes in the mean genetic gains and genetic variance of the top 1% of individuals evaluated for each trait. (**a**) Changes in the mean genetic gains of the top 1% of individuals. (**b**) Changes in the genetic variance of the top 1% of individuals. *t* represents the generation at which the second-round cross was made. Green dashed line: Scenario 1; Purple solid line: Scenario 2

## Discussion

In the present study, we evaluated the effectiveness of using UC to select cross pairs in interpopulation crosses aimed at simultaneously improving multiple traits. Additionally, to propose specific guidelines for inter-population crosses in the context of GS, we examined the optimal timing of mating. When assessing genetic gains using the selection index, Scenario 2, which employed UC, showed no noticeable difference in the mean genetic gains of the entire population compared with that in conventional selection based on GEBV; however, it significantly outperformed conventional selection in terms of the mean genetic gains of the top 1% of individuals (Fig. 3). This result can be attributed to Scenario 2, which maintained a higher genetic variance across generations. Previous studies have also indicated that maintaining high genetic variance plays a crucial role in the genetic improvement of a population (Jannink et al. 2010). In Scenario 1, individuals were selected based on the mean GEBV, which likely allowed crosses between genetically similar individuals. In contrast, in Scenario 2, cross pairs were selected based on UC, considering both the mean genotypic values and genetic variance, thereby ensuring better compatibility between parents. Moreover, the greatest genetic gains were observed when the second-round cross was performed at G_4_, likely because of the higher genetic variance maintained in that generation than in the other generations. Fig. S4 shows changes in genetic variance when inbreeding via SSD was repeated after a single cross. Genetic variance initially increased because of segregation from the cross but gradually plateaued as the population became genetically fixed. Performing crosses during generations with high genetic variance enables the selection of superior individuals, leading to greater genetic improvement. In extreme cases, performing crosses when the population is genetically fixed results in maximum genetic improvement. However, considering the balance between the costs of accelerating generations and the degree of improvement achieved through crosses, the optimal timing for crossing is likely around G_3_ and G_4_ when the increase in genetic variance starts to plateau.

Moreover, when focusing on improvement in each trait, Scenario 1, wherein GEBV was used for selection, demonstrated a superior performance than that of Scenario 2 regarding improvement in perillaldehyde content. In contrast, Scenario 2, wherein UC was utilized to select cross pairs, showed a significantly greater improvement in rosmarinic acid content (Fig. 7). When examining the final genetic gains for each trait, it was observed that perillaldehyde content exhibited minimal improvement compared with that in the initial generation; however, substantial improvements were observed in rosmarinic acid content. This result suggests that Scenario 2, which effectively facilitated the improvement in rosmarinic acid content, outperformed Scenario 1 in terms of improvement in the selection index. When comparing the GEBV of ideal individuals, those possessing favorable alleles at all markers, with the maximum GEBV of the initial population, perillaldehyde content achieved a multiple of 3.78, whereas rosmarinic acid content achieved a multiple of 5.37 (data not shown). Perillaldehyde content is regulated by a limited number of major genes with significant effects, whereas rosmarinic acid is controlled by genes with relatively small effects. The limited improvement in perillaldehyde levels can be attributed to the accumulation of favorable alleles in major genes in the initial generation in superior individuals. Conversely, improvement in rosmarinic acid content required the accumulation of favorable alleles across many markers, indicating a greater potential for improvement. We hypothesized that UC may provide more accurate predictions for traits governed by polygenes, such as rosmarinic acid, because of the assumption of a normal distribution for the GEBVs of the progeny. Allier et al. (2019) suggested that allele pyramiding is more suitable for traits controlled by a few major QTLs.

In Scenario 2, selecting cross pairs from the initial population based on the predicted genotypic value of the generation used for the second-round cross rather than that of G_15_ resulted in slightly greater genetic gains (Fig. 4), likely because UC introduces a slight prediction bias when selection is applied (Allier et al. 2019). Therefore, when the predicted genotypic value of the final generation is used as the basis, the selection of cross pairs based on the predicted genotypic value of the generation used for the second-round cross may cause a deviation in the distribution of progeny GEBVs in the final generation compared with that in the initial prediction.

In this study, the estimated marker effects were used as true marker effects in the breeding simulation, and no updates of the GP model were performed. However, it has been asserted that the prediction accuracy of the GP model is directly related to the effectiveness of selection based on UC (Müller et al. 2018). Although the advantage of selecting cross pairs using UC with model updates has been demonstrated in a study assuming recurrent selection (Sakurai et al. 2024), future research should consider the prediction accuracy of GP or heritability. Such studies will help advance our understanding of the effectiveness of inter-population crosses in the context of GS.

The interpopulation crossing is a breeding method empirically used when improving multiple traits within a single biparental population is challenging. In this study, we used two red perilla biparental populations as the material, which required the simultaneous improvement of multiple medicinal compounds. However, this method can only be applied to self-pollinating crops with more than two biparental populations and is particularly effective for crops that are yet to achieve considerable improvement in genetic traits through breeding programs, such as NUS. A few varieties of NUS often exhibit stable performance across many agronomic traits (Hunter et al. 2019).

Considering limited genetic and economic resources, breeding can first be performed in multiple biparental populations to improve specific traits within each population, followed by interpopulation crosses to improve multiple traits simultaneously. Furthermore, as each biparental population, developed independently, is expected to maintain distinct genetic variations, incorporating the effects of different alleles from each population into UC calculations is necessary. The results obtained in this study provide concrete guidelines for using interpopulation crosses to achieve the simultaneous improvement of multiple traits.

## Acknowledgments

The seeds of the cross parents of the two perilla populations used in this study were provided by the GenBank Project for Agricultural Biological Resources of the National Institute of Agrobiological Sciences. We are grateful to Ms. Terue Kurosawa for cultivating the plant materials, Dr. Yoichi Aoki for measuring the medicinal compounds of perilla, and all the technical staff at Tsumura Co., and Kazusa DNA Research Institute, Japan for their support.

## Statements and Declarations

### Author contributions

All authors contributed to the conception and design of this study. Conceptualization: KS, KH, HI. Data curation: SK. Formal analysis: SK, KS. Funding acquisition: TT, HI. Investigation: TT. Project administration: HI. Resources: TT, MS, KS, SI. Software: SK. Supervision: HI. Visualization: SK. Writing—original draft: SK. Writing—review and editing: SI, HI. All authors have read and agreed to the published version of the manuscript.

### Data availability

The datasets generated and analyzed during the current study are available in the “Sei- Kinoshita/RPSP” repository on GitHub (https://github.com/Sei-Kinoshita/RPSP). Genome assembly data and predicted gene sequences were obtained from Plant GARDEN (https://plantgarden.jp/ja/list/t48386p/). The obtained genome sequence reads are available from the DDBJ Sequence Read Archive (DRA) under the accession number DRA019597. The BioProject accession number of the submitted dataset is PRJDB19161.

### Ethics approval

Not applicable.

### Consent to participate

Not applicable.

### Consent for publication

Not applicable.

