## Supplemental File for "Optimization of crossing strategy based on the usefulness criterion in inter-population crosses considering different genetic effects among populations"

**Sei Kinoshita [1], Kengo Sakurai [1], Kosuke Hamazaki [2], Takahiro Tsusaka [3], Miki Sakurai [3], Terue Kurosawa [3], Youichi Aoki [3], Kenta Shirasawa [4], Sachiko Isobe [1], and Hiroyoshi Iwata [1,*]**

[1] Graduate School of Agricultural and Life Sciences, University of Tokyo, Tokyo, Japan

[2] RIKEN Center for Advanced Intelligence Project, Chiba, Japan

[3] TSUMURA & CO., Ibaraki, Japan

[3] Kazusa DNA Research Institute, Chiba, Japan

*** Correspondence:**
Corresponding Author


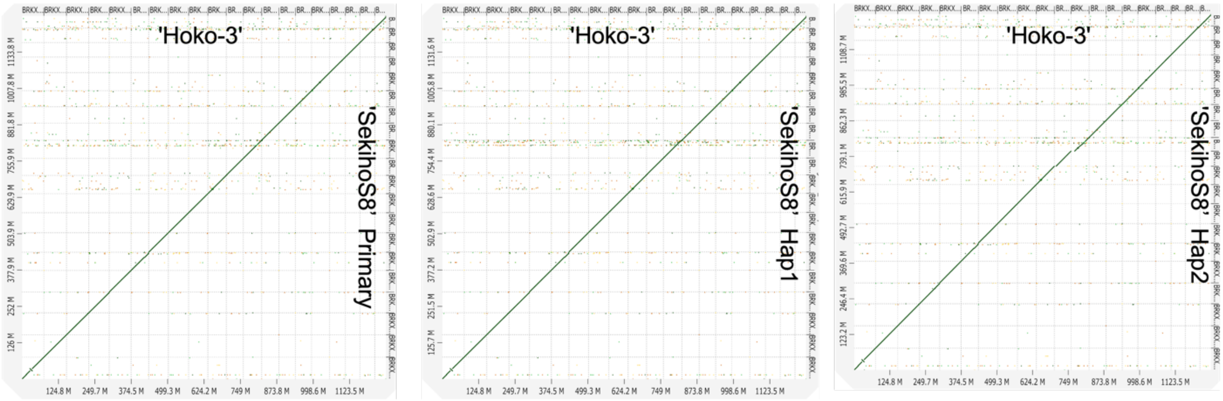


**Fig. S1** Comparison of the genome structures of 'Hoko-3' and 'SekihoS8'.

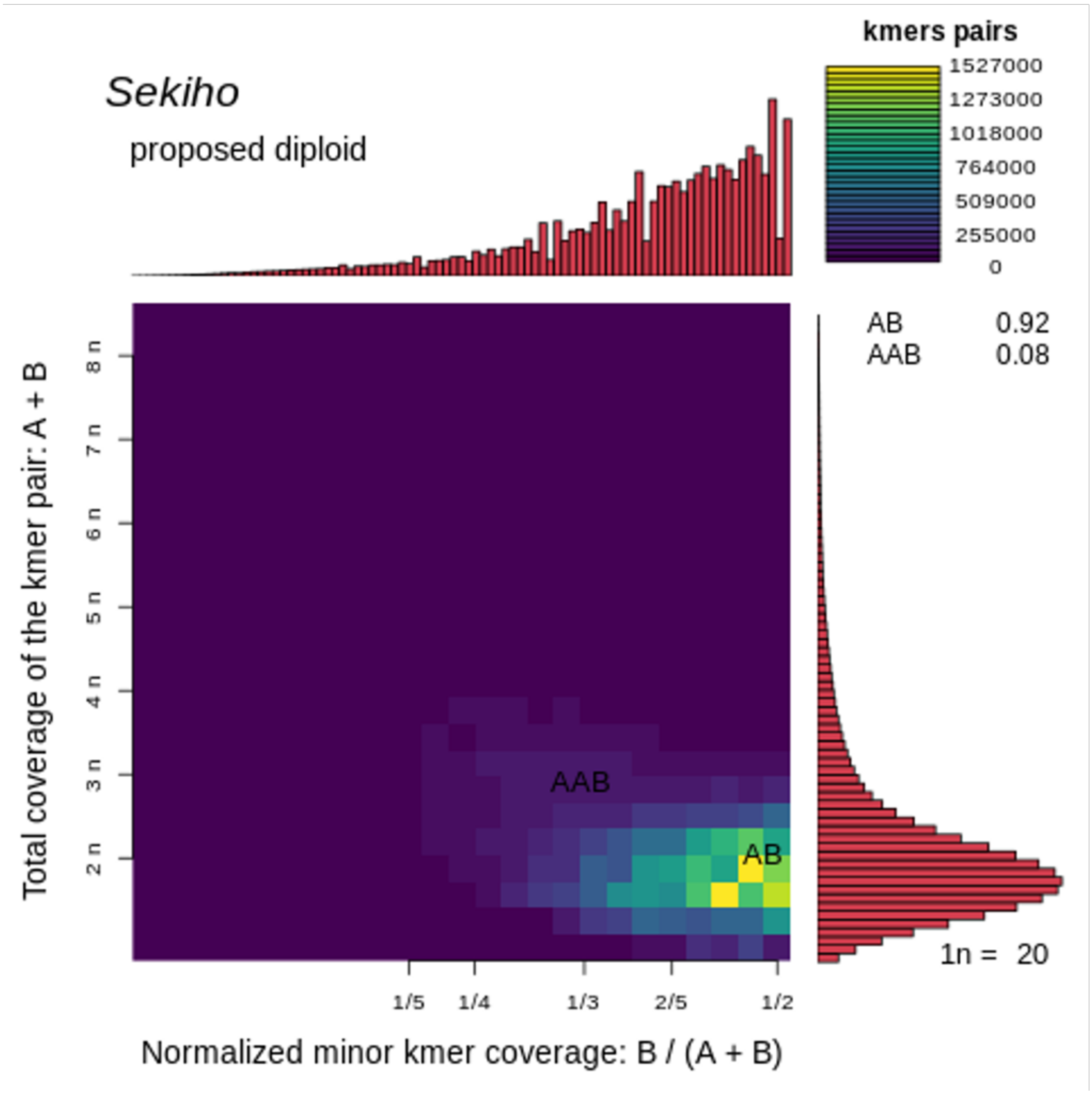


**Fig. S2** Genome structure estimation of 'SekihoS8' using Smudgeplot. The analysis was conducted using the primary assembly.

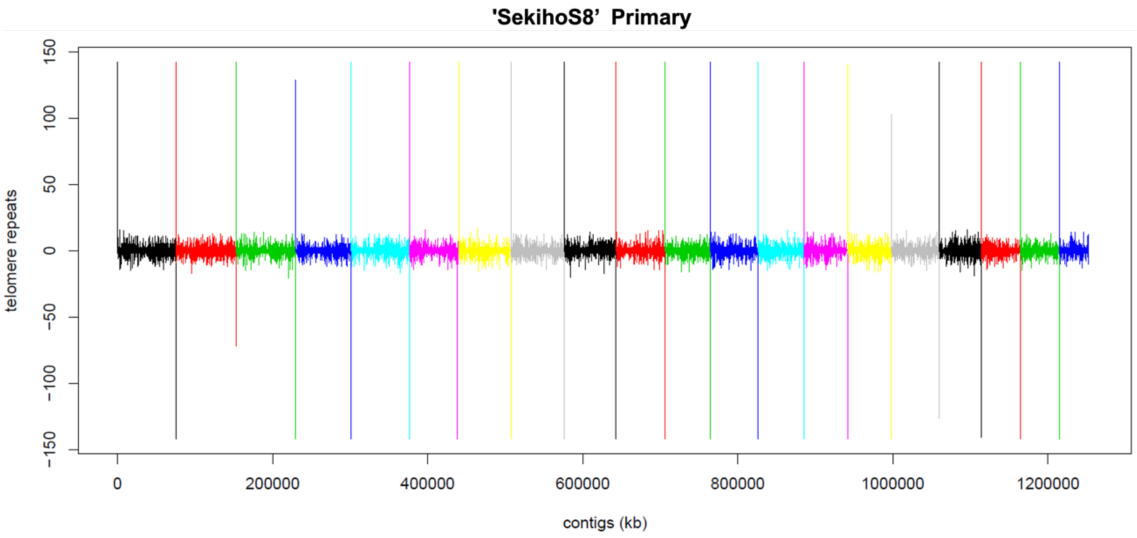

**Fig. S3A** Results of telomere sequence detection on the 'SekihoS8' primary genome

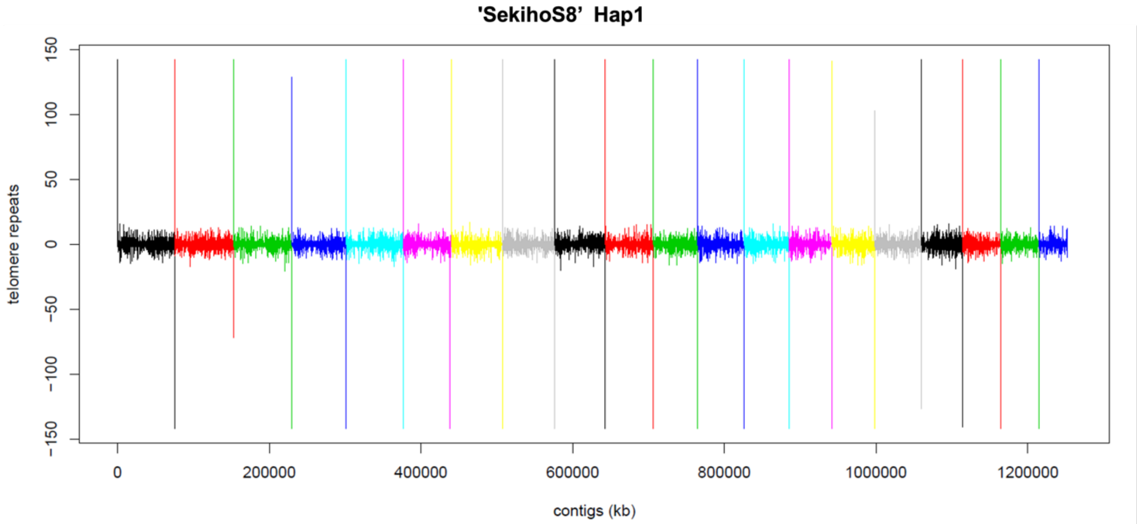

**Fig. S3B** Results of telomere sequence detection on the 'SekihoS8' Hap1 genome

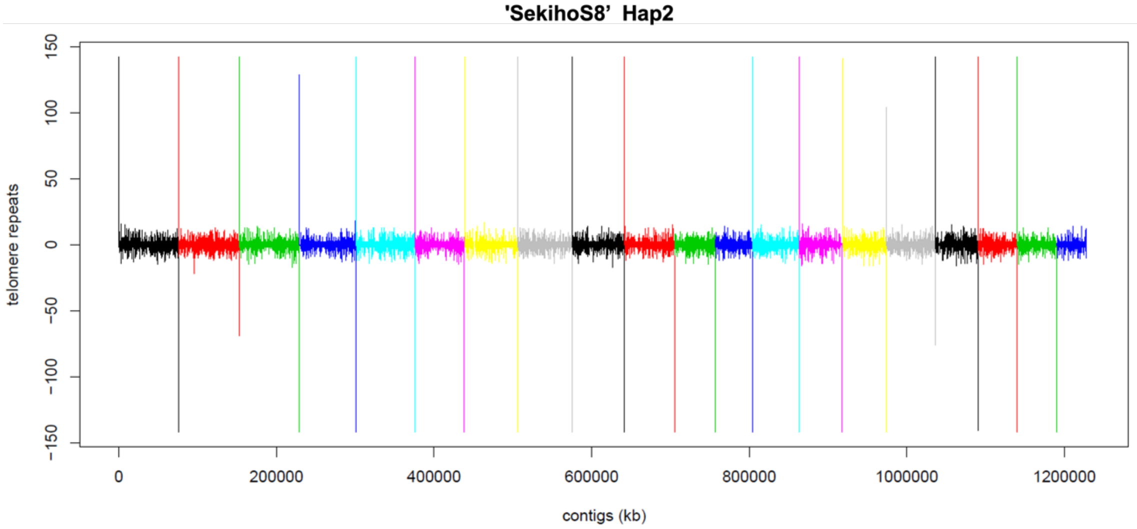


**Fig. S3C** Results of telomere sequence detection on the 'SekihoS8' Hap2 genome

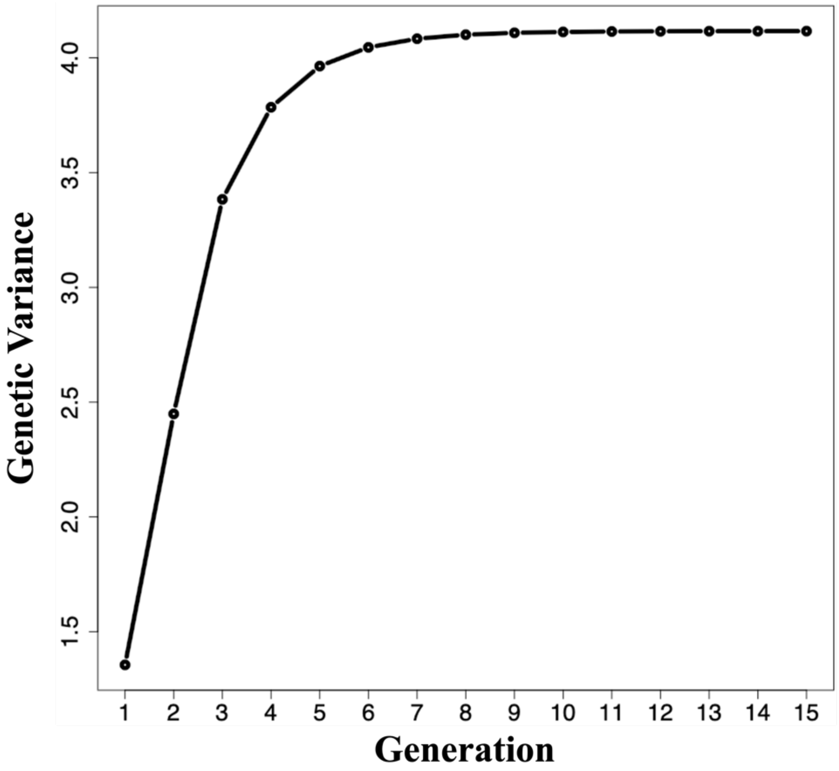


**Fig. S4** Change in genetic variance during inbreeding via single-seed descent (SSD) repeated after a single cross.

**Supplementary Table kdri1. Statistics of the assembled 'SekihoS8' genome.**

| Result of assembly by hifiasm | | | |
|---|---|---|---|
| | Hap1 | Hap2 | Primary |
| Number of sequences | 357 | 161 | 343 |
| Total length (bp) | 1,268,085,799 | 1,235,131,714 | 1,270,581,718 |
| Average length (bp) | 3,552,061 | 7,671,626 | 3,704,320 |
| Max length (bp) | 77,417,536 | 77,371,512 | 77,417,536 |
| Min length (bp) | 20,070 | 21,559 | 20,070 |
| N50 length (bp) | 55,681,988 | 55,476,749 | 55,956,609 |
| After removal of candidate chloroplast and mitochondrial genome sequences | | | |
| | Hap1 | Hap2 | Primary |
| Number of sequences | 121 | 106 | 108 |
| Total length (bp) | 1,257,287,594 | 1,231,835,063 | 1,259,745,714 |
| Average length (bp) | 10,390,807 | 11,621,086 | 11,664,312 |
| Max length (bp) | 77,417,536 | 77,371,512 | 77,417,536 |
| Min length (bp) | 21,559 | 21,559 | 21,559 |
| N50 length (bp) | 55,681,988 | 55,476,749 | 55,956,609 |
| Aligned to Pfru_yukari_1.0 using Ragoo | | | |
| | Pfru_SekihoH1_1.0 | Pfru_SekihoH2_1.0 | Pfru_SekihoUp_1.0 |
| Number of sequences | 20 | 20 | 20 |
| Total length (bp) | 1,257,297,694 | 1,231,843,663 | 1,259,754,514 |
| Average length (bp) | 62,864,885 | 61,592,183 | 62,987,726 |
| Max length (bp) | 77,417,536 | 77,371,512 | 77,417,536 |
| Min length (bp) | 42,555,208 | 41,231,751 | 42,872,164 |
| N50 length (bp) | 63,696,810 | 63,607,629 | 63,696,810 |
| BUSCO v5.2.2 (odb10, %, n = 1,614) | | | |
| | Pfru_SekihoH1_1.0 | Pfru_SekihoH2_1.0 | Pfru_SekihoUp_1.0 |
| Complete | 99.4 | 99.4 | 99.4 |
| Single | 4.2 | 9.7 | 4.2 |
| Duplicate | 95.2 | 89.7 | 95.2 |
| Fragment | 0.2 | 0.2 | 0.2 |
| Missing | 0.4 | 0.4 | 0.4 |

**Supplementary Table kdri2. Statistics of the predicted genes on the 'SekihoS8' genomes.**

| Result of assembly by hifiasm | | | |
|---|---|---|---|
| | Pfru_SekihoH1 | Pfru_SekihoH2 | Pfru_SekihoUp |
| Number of sequences | 65,078 | 62,895 | 65,161 |
| Total length (bp) | 81,205,836 | 78,622,701 | 81,312,075 |
| Average length (bp) | 1,248 | 1,250 | 1,248 |
| Max length (bp) | 16,365 | 16,368 | 16,365 |
| Min length (bp) | 102 | 102 | 102 |
| N50 length (bp) | 1,632 | 1,635 | 1,632 |
| BUSCO v5.2.2 (odb10, %, n = 1,614) | | | |
| | Pfru_SekihoH1_1.0.fasta | Pfru_SekihoH2_1.0.fasta | Pfru_SekihoUp_1.0.fasta |
| Complete | 99.1 | 98.7 | 99.1 |
| Single | 4.2 | 9.7 | 4.2 |
| Duplicate | 94.9 | 89.0 | 94.9 |
| Fragment | 0.6 | 0.7 | 0.6 |
| Missing | 0.3 | 0.6 | 0.3 |
